# Publicly available transcriptomes provide the opportunity for dual RNA-Seq meta-analysis in *Plasmodium* infection

**DOI:** 10.1101/576116

**Authors:** Parnika Mukherjee, Gaétan Burgio, Emanuel Heitlinger

**Affiliations:** Department of Molecular Parasitology, Humboldt University Berlin, Germany; Research Group Ecology and Evolution of Molecular Parasite-Host Interactions, Leibniz-Institute for Zoo and Wildlife Research (IZW), Berlin, Germany; Department of Immunology and Infectious Diseases, John Curtin School of Medical Research, Australian National University, Canberra, ACT, Australia

## Abstract

Dual RNA-Seq is the simultaneous transcriptomic analysis of interacting symbionts, for example, in malaria. Potential cross-species interactions identified by correlated gene expression might highlight interlinked signaling, metabolic or gene regulatory pathways in addition to physically interacting proteins. Often, malaria studies address one of the interacting organisms – host or parasite – rendering the other “contamination”. Here we perform a meta-analysis using such studies for cross-species expression analysis.

We screened experiments for gene expression from host and *Plasmodium*. Out of 171 studies in *Homo sapiens, Macaca mulatta* and *Mus musculus*, we identified 63 potential studies containing host and parasite data. While 16 studies (1950 samples) explicitly performed dual RNA-Seq, 47 (1398 samples) originally focused on one organism. We found 915 experimental replicates from 20 blood studies to be suitable for co-expression analysis and used orthologs for meta-analysis across different host-parasite systems. Centrality metrics from the derived gene expression networks correlated with gene essentiality in the parasites. We found indications of host immune response to elements of the *Plasmodium* protein degradation system, an antimalarial drug target. We identified well-studied immune responses in the host with our co-expression networks as our approach recovers known broad processes interlinked between hosts and parasites in addition to individual host and parasite protein associations.

The set of core interactions represents commonalities between human malaria and its model systems for prioritization in laboratory experiments. Our approach might also allow insights into the transferability of model systems for different pathways in malaria studies.

**Importance:** Malaria still causes about 400,000 deaths a year and is one the most studied infectious diseases. The disease is studied in mice and monkeys as lab models to derive potential therapeutic intervention in human malaria. Interactions between *Plasmodium* spp. and its hosts are either conserved across different host-parasite systems or idiosyncratic to those systems. Here we use correlation of gene expression from different RNA-Seq studies to infer common host-parasite interactions across human, mouse and monkey studies. We, firstly, find a set of very conserved interactors, worth further scrutiny in focussed laboratory experiments. Secondly, this work might help assess to which extent experiments and knowledge on different pathways can be transferred from models to humans for potential therapy.

## Introduction

Transcriptomes are often analysed in a first attempt to understand molecular, cellular and organismic events. A comprehensive profile of RNA expression can be obtained using high-throughput sequencing of cDNA from reverse transcripted expressed RNA. Such RNA-sequencing (RNA-Seq) provides high technical accuracy at a reasonable cost, making it the current method of choice for transcriptomics [1,2].

In an infection experiment, RNA-Seq can assess host and pathogen transcriptomes simultaneously if RNA from both organisms is contained in a sample. On the one hand, it has been proposed to analyse transcriptomes of both organisms involved in an infection for a more complete understanding of the disease [3]–[5], such as virulence of a pathogen resulting from interlinked processes of both host and pathogen (host-pathogen interactions). This approach is called dual RNA-Seq. Some recent studies on malaria pathogenesis make use of dual RNA-Seq to study the host and the parasite simultaneously. On the other hand, researchers intending to study one of the two organisms, the target, might consider transcripts from the non-target organism, “contamination”. Malaria research is indeed traditionally designed to target one organism, either the host or the parasite. Nevertheless, expression of “contaminant” transcripts potentially corresponds to response to stimuli.

Malaria is the most thoroughly investigated disease caused by a eukaryotic organism and the accumulation of these two types of studies, RNA-Seq with “contaminants” and intentional dual RNA-Seq, provides a rich resource for meta-analysis. In case of malaria, unlike in bacterial or viral infections, both the parasite, *Plasmodium* spp. and the host are eukaryotic organisms with similar transcriptomes [4]. Their messenger RNA (mRNA) have a long poly-A tail at the 3’-end, therefore host and parasite mRNA is selected simultaneously when poly-dT priming is used to amplify polyadenylated mRNA transcripts [4,5]. This makes most malaria transcriptome datasets potentially suitable for dual RNA-Seq analysis.

Such a meta-analysis can use co-regulated gene expression to infer host-parasite interactions. In the mammalian intermediate host, *Plasmodium* spp. first multiply in the liver and then invade red blood cells (RBCs) for development and asexual expansion. While the nuclear machinery from both host and parasite cells produces mRNA in the liver, RBCs are enucleated and transcriptionally inactive in the mammalian host. In the blood stage of the infection, leukocytes or White Blood Cells (WBCs) provide most of the transcriptomic response and are, thus, the source of host mRNA. Correlation of mRNA expression can be indicative of different types of biological interactions: protein products could be directly involved in the formation of complexes and might therefore be produced at quantities varying similarly under altered conditions. Alternatively, involvement in the same biological pathways can result in co-regulated gene expression without physical interactions. This broad concept of interaction has long been exploited in single organisms (e.g. [6]-[8]). We (and others before [9,10]) propose to extrapolate this to interactions between the host and the parasite. It can be expected that a stimulus presented by the parasite to a host causes host immune response and the parasite in turn tries to evade this response, creating a cascade of genes co-regulated at different experimental conditions.

Here we showed that existing raw sequencing read datasets collectively present the potential to answer questions that have not been investigated by individual studies on malaria. We explored this potential by conducting a comparative meta-analysis of dual RNA-Seq transcriptomes of *Plasmodium* and its hosts in the blood stage of infection. Since rodent and simian malaria are often used as laboratory models for human malaria, we demonstrated the availability and suitability of mRNA sequencing data from three evolutionarily close hosts - *Homo sapiens, Macaca mulatta* and *Mus musculus* - and their respective *Plasmodium* species. We summarised available data and conceived an approach to elucidate host-parasite interactions using orthologs across different host-parasite pairs. We demonstrated that this approach provides meaningful results by cross-validation of information content from networks from different host-parasite systems. We found that these networks highlight known functional determinants of host-parasite interactions in broad categories such as invasion and calcium ion homeostasis for the parasite and host immune response and cadherin mediated cell-cell adhesion on the host side. We also found novel interactions at a finer scale potentially worth further investigation.

## Methods

### Data review and curation of potentially suitable studies

Sequence data generated in biological experiments is submitted to one of the three mirroring databases of the International Nucleotide Sequence Database Collaboration (INSDC): NCBI Sequence Read Archive (SRA) [11], EBI European Nucleotide Archive (ENA) [12] and DDBJ Sequence Read Archive (DRA) [13]. We used SRAdb v1.36.0 [14], a Bioconductor/R package [15], to query SRA [11] for malaria RNA-Seq studies with the potential to provide host and *Plasmodium* reads for our meta-analysis. We first selected studies with library_strategy “RNA-Seq” and with “Plasmodium” in the study title, abstract or sample attributes fields using the function dbGetQuery(). Then we used the getSRA() function with the query “(malaria OR Plasmodium) AND RNA-Seq”. This function searches all fields. We manually curated the combined results and added studies based on a literature review using the terms described for the getSRA() function in PubMed and Google Scholar. We disregarded studies on vectors, non-target hosts and those derived from cell culture conditions devoid of transcriptionally active host cells. In these databases, all experiments submitted under a single accession are given a single “study accession number” and are collectively referred to as a “study” here onwards. We used prefetch and fastq-dump functions from SRAtoolkit v2.8.2, to download all replicate samples (called runs in the databases) of the selected studies. The curation and the download of the studies was performed on 21 January, 2019 and updated on 24 July, 2020.

### Mapping and quantification of gene expression

We mapped sequencing reads onto concatenated host and parasite reference genomes using STAR v2.6.0c [16,17] with default parameters. Only runs with at least 70% uniquely mapping reads were considered for further analysis. Host and *Plasmodium* sp. genome assemblies and gene annotation files were downloaded from Ensembl version 43. Simultaneous mapping against both genomes should avoid non-specific mapping of reads in regions conserved between host and parasites. We quantified the sequencing reads mapped to exons using the countOverlaps function of the GenomicRanges package v3.7 [18].

### Identification of co-expressed genes via correlation techniques

Reid and Berriman [9] recommended using empirical p-values for the analysis of gene co-expression. This allows not only to scrutinise housekeeping genes likely to show almost uniform expression under different experimental conditions but also requires fewer assumptions about the quality of the input data. We computed correlation indices for each gene-pair and obtained empirical p-values by comparison against null distributions computed using permutations of the given data, instead of assuming a theoretical null distribution. This is a robust way to estimate whether gene pairs are correlated because of specific events (treatment condition, time point) and not by chance (e.g., housekeeping genes) [19,20]. To obtain p-values corrected for multiple testing, since host and parasite genomes total at nearly 30,000 genes, the number of permutations would have to be around 10^12^ for a resolution of 0.1% FDR. As computational costs for these permutations are too high we used uncorrected p-values to rank genes as initially proposed by Reid and Berriman [9]. Here we only consider uncorrected p-values of 10^−5^ (in 100,000 permutations) as evidence of co-regulation.

### Selection of runs for analysis

The construction of host-parasite gene co-expression networks require RNA-Seq runs to have both host and parasite transcript expression. To fulfil these criteria, we implemented thresholds based on host and parasite transcript expression, on the selection of runs. If a study had at least 5 runs with 50% detectable transcriptome expression from the host and from the parasite, we defined 2 thresholds for runs: 1) “intermediate” (int) with at least 50% detectable host and parasite transcriptome expression and 2) “stringent” (str) with 70% detectable host and parasite transcriptome expression. If a single study had less than 5 runs in the intermediate threshold, these runs were pooled with runs from other such studies.

To compare sub-datasets with runs selected at different thresholds and including all runs without thresholds, we calculated the Jaccard index for every pairwise combination of these datasets.

The Jaccard Index is defined as 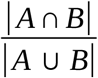 [21], where A and B are the set of bipartite edges in from two datasets. To include the maximum amount of host and parasite data and to use the best dataset from each study, we calculated the sum of all Jaccard Indices from each dataset. We chose the sub-dataset with the highest Jaccard index for a given study for further analysis and concatenated these representative studies into an “overall” dataset to compute possible conserved interactions from all host-parasite systems.

### Identification of orthologs

Orthologs are genes in different species that have been derived from a single gene in the last common ancestor of those species by vertical descent. This means that orthologous genes from different species, in general, are likely to perform the same function [22].

Whole genome based predictions of the proteome for host species were downloaded from Ensembl release 43 and for *Plasmodium* from PlasmoDB release 37. Orthologs were identified using OrthoFinder v2.2.7 [23] wherein blastp [24] results for all-versus-all protein comparisons are clustered using the MCL clustering algorithm [25] (tools bear default parameters for OrthoFinder). One-to-one ortholog clusters, or orthogroups, were recovered for hosts and for parasites.

### Network and functional analyses

Bipartite networks/graphs are graphs in which the nodes of the graph can be divided into two independent sets, and the nodes from one set connect only with nodes in the other set [26,27]. To visualise host-parasite interactions, bipartite networks were constructed with R package igraph v1.2.5 [28] and Cytoscape v3.8.0 [29] where the nodes were the genes and their interaction were represented with an edge. To cluster nodes into network modules, we used the edge-betweenness algorithm (function edge_betweenness()) from igraph. The UpSetR package (v1.4.0) [30] in R was used to visualise intersection sizes of overlapping edges between networks.

For functional analysis of the genes found in host-parasite interactions from co-expression analysis, their Gene Ontology [31] terms were found using the package topGO v2.36.0 and enriched using Kolmogorov-Smirnov test [32]. PlasmoDB [33] and Malaria.tools [34] databases were used for looking up gene functions and stage-associated transcript expression.

### Statistical modeling

Using the R package fitdistrplus v1.1-1, we determined how Relative Growth Rate (RGR) [35] and Mutagenesis Index Score (MIS) [36] are distributed. We used the betareg package in R to model RGR and MIS using centrality properties of our networks.

The centrality properties measured for our networks were node degree (DG), eigenvector centrality (EC) and betweenness (BW) using their dedicated functions in igraph package in R: “degree()”, “betweenness()” and “eigen_centrality()”. RGR and MIS (as response variables) were modelled first with a single centrality measure as a predictor and then with combinations of two centrality measures: DG with EC and DG with BW. “Nested” models were compared based on likelihood-ratio tests and the more complex model was accepted when delta likelihood exceeded 2. In addition to this, the Akaike Information Criterion (AIC) was computed for each model to compare models and again models were considered differing in explanatory power at a delta-AIC of 2. Models on different datasets (and different response variables) were compared without explicit statistical testing discussing differences in p-values for variables in question. In general, R (v3.4.3 - 4.0.0) [37] was used for analysis. All code is available at https://doi.org/10.5281/zenodo.4535898 (as used in this manuscript) and github repository https://github.com/parnika91/CompBio-Dual-RNAseq-Malaria (under further development).

## Results

### Potentially suitable studies for human, mouse and simian malaria

We found 63 potentially suitable studies [3,27,38-97] (Supplementary Table S1) on querying SRA and performing a literature review. The host organism for 27 studies was *Homo sapiens*, for 26, *Mus musculus* and for 10, *Macaca mulatta*. The infecting parasites were *P. falciparum, P. vivax* and *P. berghei* in human studies (including artificial infections of human hepatocyte culture with *P. berghei*), *P. yoelii, P. chabaudi* and *P. berghei* in mouse studies and *P. cynomolgi* and *P. coatneyi* in macaque studies (Table 1). For 16 out of the 63 studies, the authors state that they intended to simultaneously study host and parasite transcriptomes. This includes 8 studies from MaHPIC (Malaria Host-Pathogen Interaction Center), based at Emory University, which made extensive omics measurements in simian malaria. The original focus of the remaining 47 studies was on the parasite in 23 and on the host in 24 cases. Our collection of studies comprises data derived from blood and liver for all three host organisms human, mouse and macaque. Experiments performed on mouse blood focus on the parasite instead of the host (12 vs. 0). Studies on human blood infection focus more often on the host immune response than on the parasite (10 vs. 6). Liver and spleen studies focus on host and parasite almost equally as often, with sources for host tissue in this case being either mice (in vivo) or hepatoma cultures (in vitro).

**Table 1.** Potentially available host-parasite systems, number of studies and runs in different tissues after database querying. Numbers are arranged as the number of studies / number of runs.

|  | Blood | Spleen | Liver | Brain | Bone marrow | Lungs |
| --- | --- | --- | --- | --- | --- | --- |
| Human - *P. falciparum* | 14 / 855 | 0 / 0 | 0 / 0 | 0 / 0 | 0 / 0 | 0 / 0 |
| Human - *P. vivax* | 6 / 141 | 0 / 0 | 1 / 4 | 0 / 0 | 0 / 0 | 0 / 0 |
| Human - *P. berghei* | 0 / 0 | 0 / 0 | 6 / 145 | 0 / 0 | 0 / 0 | 0 / 0 |
| Mouse - *P. yoelii* | 3 / 19 | 2 / 10 | 3 / 20 | 0 / 0 | 1 / 4 | 1 / 6 |
| Mouse - *P. chabaudi* | 4 / 204 | 5 / 795 | 2 / 46 | 0 / 0 | 0 / 0 | 0 / 0 |
| Mouse - *P. berghei* | 3 / 44 | 0 / 0 | 0 / 0 | 3 / 83 | 0 / 0 | 0 / 0 |
| Monkey - *P. cynomolgi* | 5 / 811 | 0 / 0 | 2 / 59 | 0 / 0 | 0 / 0 | 0 / 0 |
| Monkey - *P. coatneyi* | 1 / 35 | 0 / 0 | 0 / 0 | 0 / 0 | 2 / 66 | 0 / 0 |

We note that 21 of the 63 studies depleted or enriched specific classes of cells from their samples. Seventeen blood-stage studies depleted or enriched host WBCs to focus expression analysis on *Plasmodium* or host immunity, respectively. Assuming this depletion was imperfect, we tested whether such samples present mRNA from both organisms. The physiologically asymptomatic liver stage [72] is an appealing target for sporozoite-derived vaccines [98]-[100], but low parasite numbers make it difficult to study *Plasmodium* transcriptomes in liver. To reduce overwhelming host RNA levels, 10 out of the 14 liver studies sorted infected hepatoma cells from uninfected cells. Three out of the remaining 4 studies were among the ones interested in the host expression during the infection and 1 enriched for host cells.

### Blood and liver samples from different studies and host parasite systems are potentially suitable for dual RNA-Seq analysis

A sample or run must provide sufficient gene expression from both host and parasite to be suitable for dual RNA-Seq co-expression analysis. Analysing the proportions of sequencing reads mapping to the host and parasite transcriptomes, we found that the original focus of the study was not always reflected (Fig 1). In many native samples (not enriched or depleted for host material), the number of host reads was found to be overwhelming. However, probably when parasitemia was very high, parasite transcriptomes were still recovered. Some examples are runs in the studies SRP032775, SRP029990 and ERP106769. Similarly, many studies using depletion or enrichment of a certain cell type prior to RNA sequencing show considerable expression of the non-target organism (enriched/depleted ‘E’ in Fig 1A). Examples are the studies ERP023892, ERP002273, ERP004598, ERP005730, ERP110375 and SRP112213. Studies depleting whole blood from leukocytes to focus on parasite transcriptomes still show considerable host gene expression and provide potentially suitable runs for the analysis of blood infection at lower intensities. Although, we note that this might come with the caveat that host expression might be biased by unequal depletion of particular cell types.

**Figure 1.**
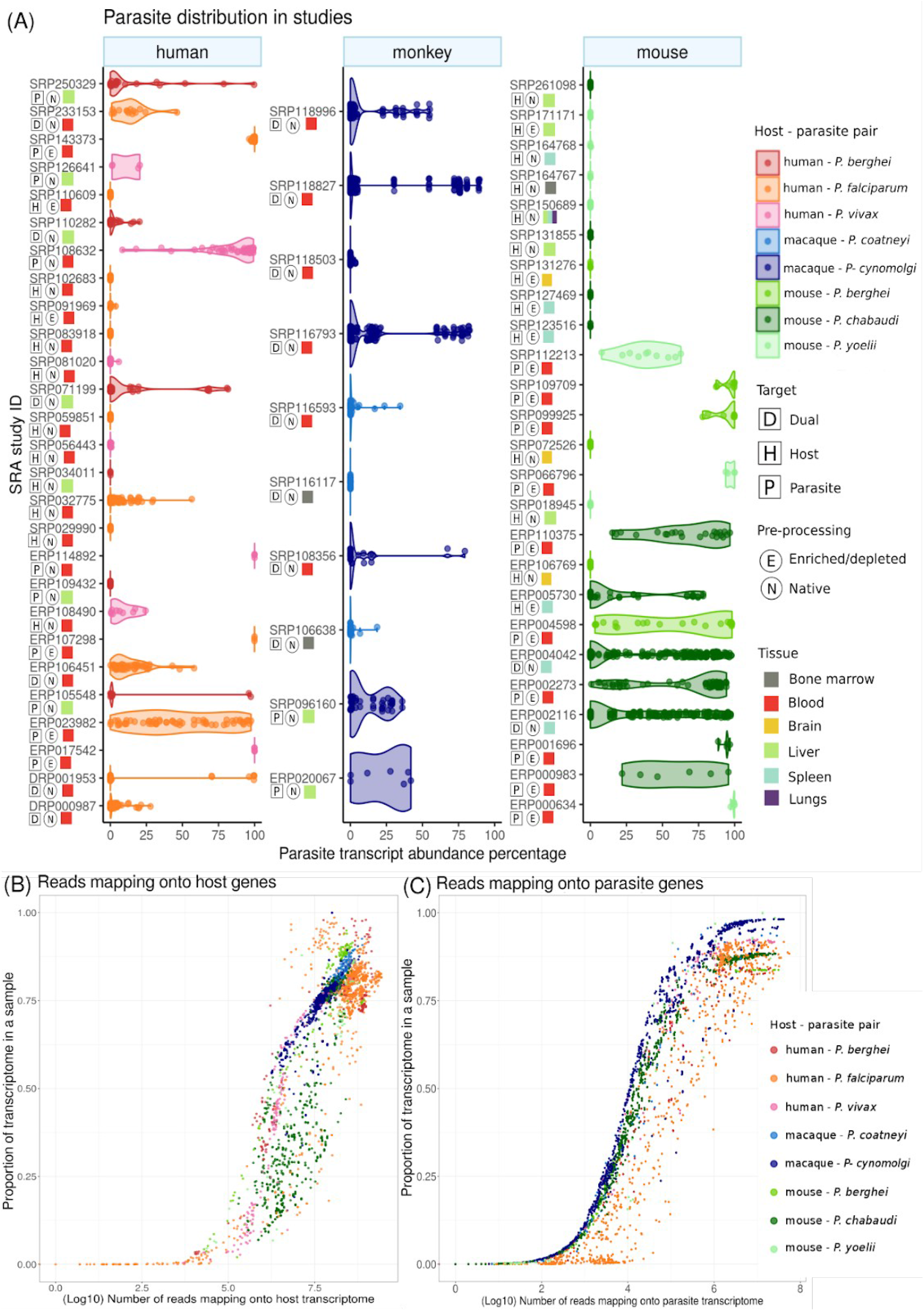
Proportion and number of sequencing reads and expressed genes from parasite and host in selected malaria RNA-Seq studies. We mapped sequencing reads from studies selected for their potential to provide both host and parasite gene expression data (total studies=63, total runs=3351) against appropriate host and parasite genomes. (A) The percentage of parasite reads (x-axis) is plotted for runs in each study (host and parasite add to 100%). The studies are categorised according to the host organisms studied and “enriched/depleted” to indicate enrichment of infected hepatocytes or depletion of leukocytes from blood. Studies labeled “dual” were originally intended to simultaneously assess host and parasite transcriptomes. We also plot the number of reads mapped against the number of expressed genes for host (B) and parasite (C). The proportion of transcriptome detected as expressed increases with sequencing depth towards the maximum of all expressed in the transcriptome.

The number of sequencing reads mapping onto a transcriptome is a major determinant of the proportion of the organism’s transcriptome detected as expressed (transcripts with at least one read mapped; Fig 1B and C). As expected, only runs that were sequenced deeply were able to capture the expression of a high proportion of the transcriptome. For both host and parasite, this plateaus at the total number of genes expressed in the transcriptome of the respective tissue in these organisms. Runs with an acceptable proportion of the transcriptome detected as expressed were chosen in our analysis of blood stage malaria below. We propose that thresholds based on the proportion of the transcriptome detected as expressed are most suitable for this selection of runs. Table 2 provides an overview of the number of studies and runs available for each of these analyses.

**Table 2.**
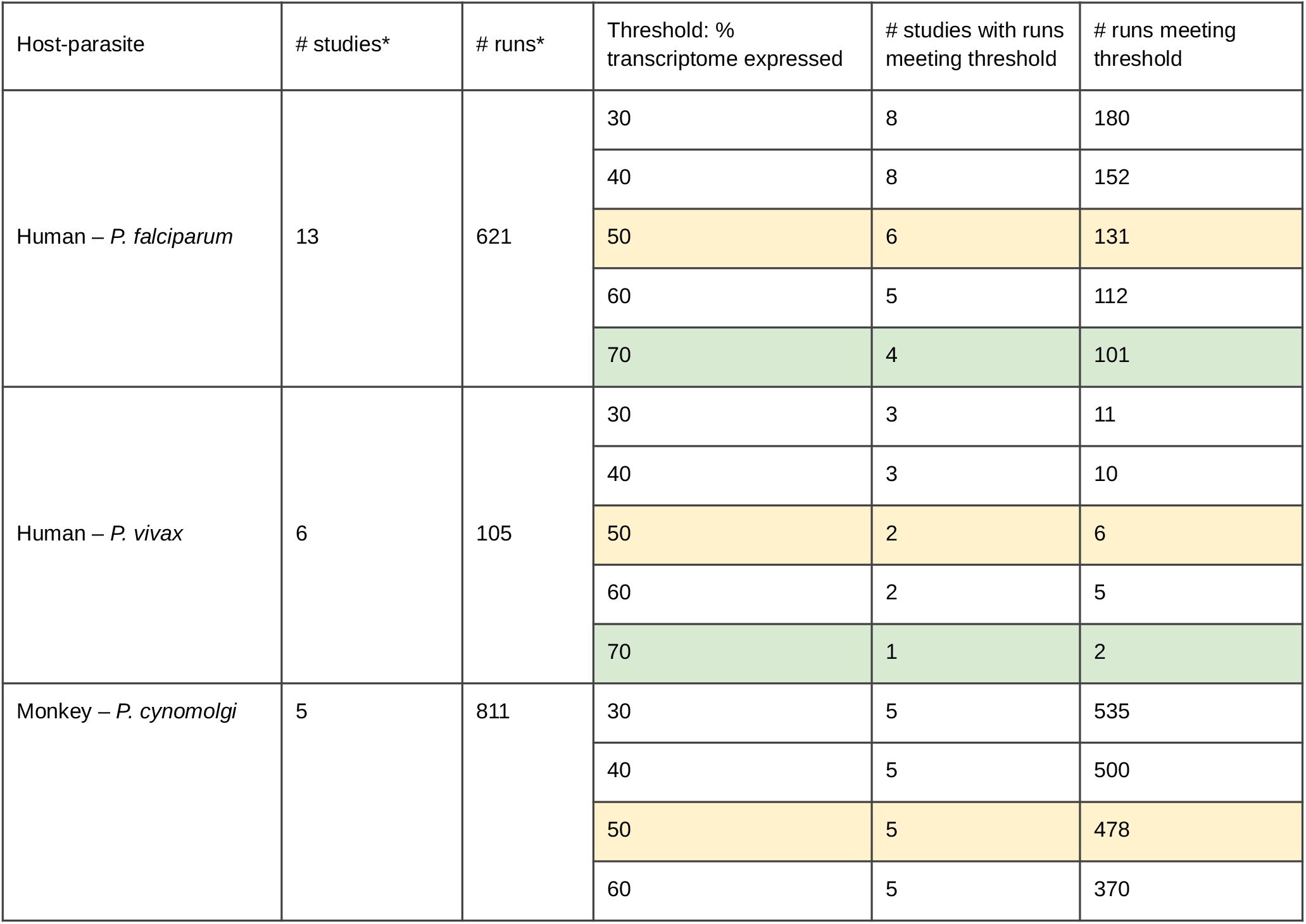
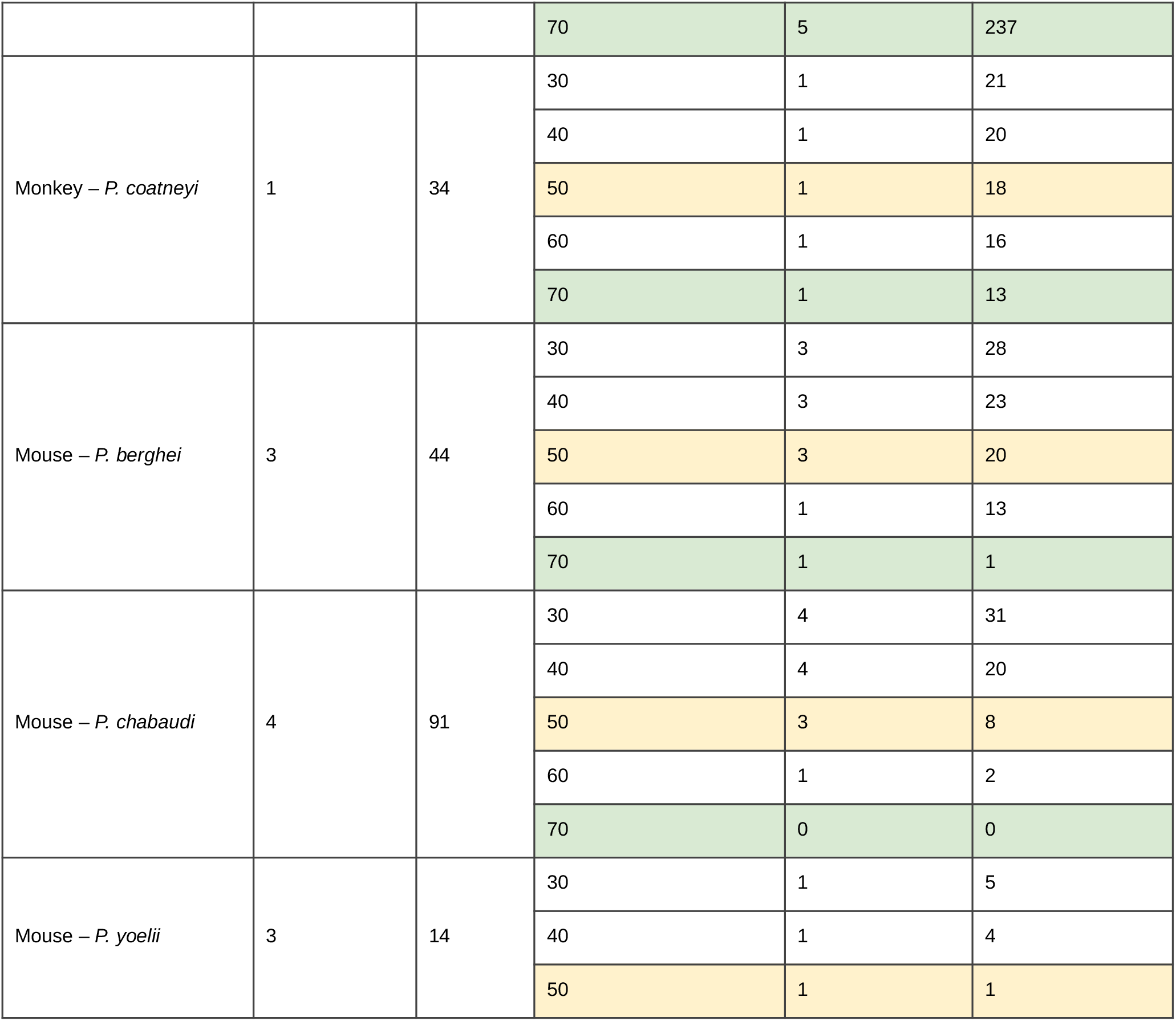
Number of blood studies for each host-parasite pair and suitability analysis of their experimental replicates (runs). [* = after applying a quality threshold of a unique read mapping proportion of 70%, yellow: intermediate threshold, green: stringent threshold]

### Evaluation of thresholds on transcriptome representation improve the analysis of co-regulated gene-expression

Deciding on the stringency of the thresholds applied to a dual RNA-Seq meta-analysis requires additional analysis. Previous studies have reported that estimated proportions of the transcriptionally active parts of the *P. falciparum* genome intraerythrocytic stages range from 60-90% [101]-[103]. Here we test different thresholds that maximise the signal common between blood studies - the number of edges shared between correlation networks for different studies (Fig 2).

**Figure 2.**
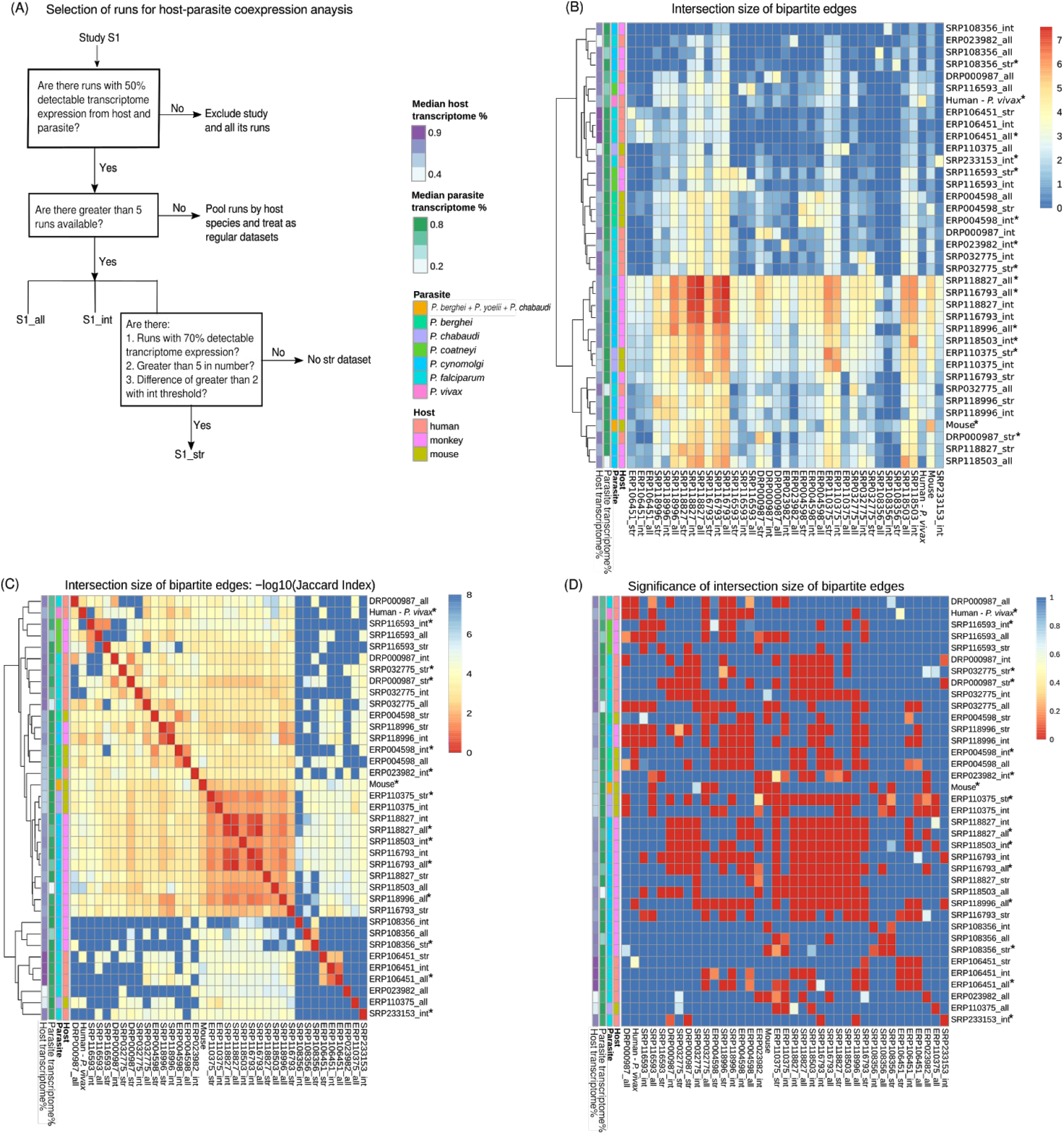
Overlapping bipartite edges between datasets across host-parasite systems. Based on the proportion of transcriptome expressed in each study, three thresholds were implemented on the selection of runs for co-expression analysis. Without thresholds “all”, includes all runs. The intermediate (“int”) and stringent (“str”) thresholds, include runs in which 50% (“int”) and 70% (“str”) of the transcriptome of both host and the parasite are detected as expressed (covered), respectively. (A) Schematic to show how samples were selected for coexpression analysis. (B) For each dataset, the median host and parasite transcriptome coverage is indicated. The heatmap shows the log10 transformed number of bipartite edges for each study on the diagonal. The set size of the overlapping bipartite edges between corresponding datasets are displayed in the remaining fields. The datasets are clustered based on euclidean distances between these set sizes. The size of the overlapping set of bipartite edges is not determined by the median parasite or host transcriptome coverage, even studies with low coverage of the parasite transcriptome provide datasets leading to high number of bipartite edges found in common with other datasets. The optimal overlap set size does not suggest a single threshold for the selection of sequencing runs - all three thresholds are found to have a high number of overlapping edges with other datasets. (C) Jaccard indices (size of intersection as a ratio of the size of union) for each dataset pair are displayed as (-)log10 transformed indices. The versions of the datasets used for further analysis are marked with an asterix. (D) The size of the intersection was tested for significance using Fisher’s exact test. The heatmap shows that all datasets led to correlation networks that overlapped with other networks from other datasets more than expected by chance. We can conclude that they contain biological signals suitable for combined analysis.

36 studies investigated the blood stages of *Plasmodium*. 13 of them provided more than 5 runs at all thresholds (schematic in Fig 2A) and were analysed as independent datasets. Similarly, 11 studies could be analysed separately with selection of runs at a “stringent” threshold of 70% transcriptome coverage. To make use of runs not meeting thresholds from studies that couldn’t be analysed separately, 6 runs from 2 studies were pooled into a combined “humanPvivax” dataset and 10 runs from 5 studies into a combined mouse dataset at intermediate thresholds. The combined mouse dataset comprised 4 runs from 2 *P. berghei* studies, 5 from 2 *P. chabaudi* studies and 1 from a *P. yoelii* study. 16 studies did not provide any runs meeting the thresholds and were therefore excluded from the analysis. Based on the sum of Jaccard indices (see Methods) of the datasets, we selected a total of 15 sub-datasets maximizing overlap between individual study networks. We concatenated them to construct the “overall” dataset: 4 studies without thresholds (“all”), 7 at intermediate (50%; “int”) and 4 at stringent (70% “str”) thresholds (marked with an asterix in Fig 2B,C,D). This means that a total of 915 runs were included in the “overall” dataset.

We can conclude from this analysis that the edges shared between different studies within and among host-parasite systems (see below for an exception regarding human-*P. falciparum* studies) outnumber random expectations and are highly significantly (p<0.001, Fisher’s exact test; between all selected studies), indicating a biological signal shared between datasets (Fig 2D). We suggest analysing the optimal version (selection of runs) of each study instead of setting a fixed (e.g. intermediate) threshold. Fig 2 indicates which different thresholds we deem most suitable for different studies and our selection for downstream analyses.

### Across different studies, across different host-parasite systems

Knowledge of one-to-one orthologs between different hosts and different parasite species can be used in the next steps to integrate across different host-parasite systems (Fig 3). Humans share 18179 1:1 orthologous genes with macaques and 17089 with mice. 13986 genes are 1:1:1 orthologs among the three host species. Similarly, 4005 one-to-one groups of orthologous genes were found among the *Plasmodium* species (Fig 3A).

**Figure 3.**
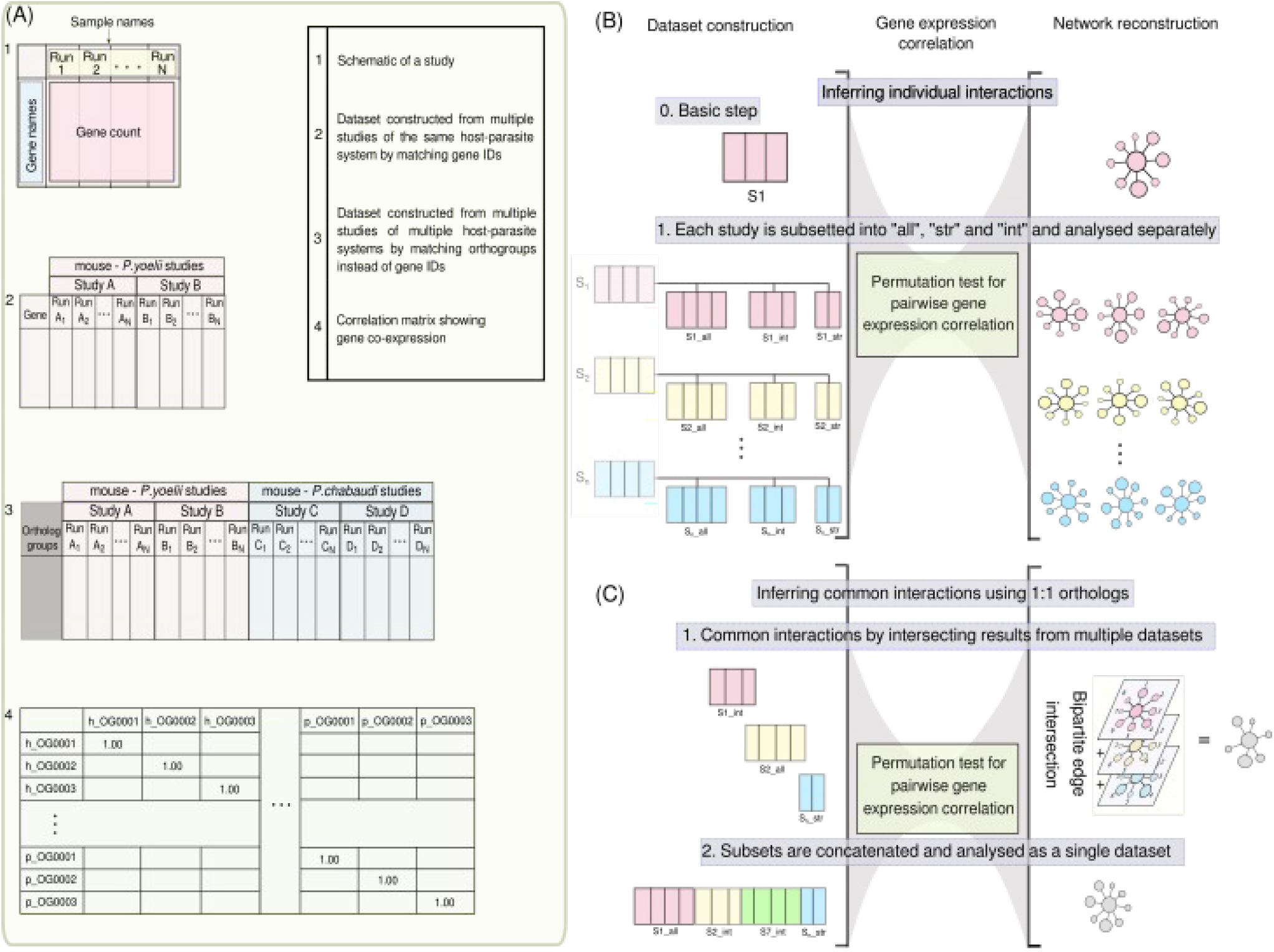
Alternative and interlinked strategies to reconstruct host-parasite interaction networks. We designed two interlinked approaches to obtain a consensus network involving multiple host-parasite systems. (A) An implementation of permutation tests derives “empirical p-values” to test correlation coefficients for significance and infers interactions. (B) The methodology used to compare transcriptome coverage thresholds (50%, intermediate, “int”; 70%, stringent, “str”) is using different subsets of runs from each individual study (“sub-datasets” S1, S2 and S3 in this illustration). As in fig 2, without thresholds “all”, includes all runs. Each subset is analysed separately to infer interactions, allowing the subset (and threshold) leading to highest information overlap with other studies to be selected. (C) shows the inference of common interactions across host-parasite systems using the knowledge of single-copy orthologs across the hosts and the parasites. Here, we can take two approaches: the first approach uses intersections in a multilayer network approach to come to a consensus of common interactions. In the second approach, we concatenate runs and correlate gene expression on those as a single “overall” dataset. In the analysis for our final results we combined these approaches mapping networks from each study onto the common network from an “overall” dataset.

We have identified two different but interlinked workflows to reconstruct a consensus network of expression correlation. A first approach integrates data from different studies of one host-parasite system by simply appending expression profiles of their runs. Expression data across host-parasite systems is combined in orthogroups and correlations of gene expression are computed for all samples simultaneously. We refer to the results of this approach as the “overall” network (Fig 3C). It has the benefits that it uses and weighs all the information sources in one concise process.

Alternatively, we search for consensus networks by comparison of individual networks from different studies (Fig 3C). Similar to the approach of optimising expression thresholds (Fig 2, 3B), we compare overlapping edges in a multilayer network analysis. Sets of overlapping edges from multiple studies and host-parasite systems offer more control when querying for similar correlation in different layers representing different host-parasite systems. Similarly, correlations from different types of tissues (blood and liver) could be combined as multilayer networks in future work.

Correlating expression on the “overall” dataset resulted in a larger network (3.64 million edges, 12652 host genes, 3996 parasite genes) than on 13 out of 15 individual study datasets - two macaque datasets were exceptions (Fig 4). 528,883 bipartite edges are found only in the “overall” dataset and not in any individual dataset. 13 - 43% of edges from the “overall” dataset are recovered in each individual dataset (median 18.5%), indicating a substantial contribution from each individual dataset to the “overall” network. The most dominant study (presenting 43% of the edges in the “overall” network in it’s own network) was a mouse study (study ID ERP004598 [91]) in which parasite gene expression was the primary focus. 91% of this study’s bipartite edges are shared with at least one other individual study in addition to the “overall” network. The second most influential study was a dual RNA-Seq study (on monkey, study ID SRP118827 [27]) that shares 89% of its bipartite edges with at least one other study (Fig 4A). This not only illustrates the contributions of studies that initially focused on one organism (“with contaminants”) in dual RNA-Seq analysis, but also that information can be pooled across different studies and host-parasite systems with our approach. Surprisingly, there were no common edges found between two human studies ([38] on Thai patients and [3] on Gambian patients, both infected with *P. falciparum*), possibly indicating differences in sampling from patients. Alternatively, this might hint towards idiosyncrasies in human blood studies using native samples with high parasitemia. The potential difficulty to apply RNA-Seq meta-analysis to these human samples highlights the need to transfer such data across model systems while controlling for concordance with the human system.

**Figure 4.**
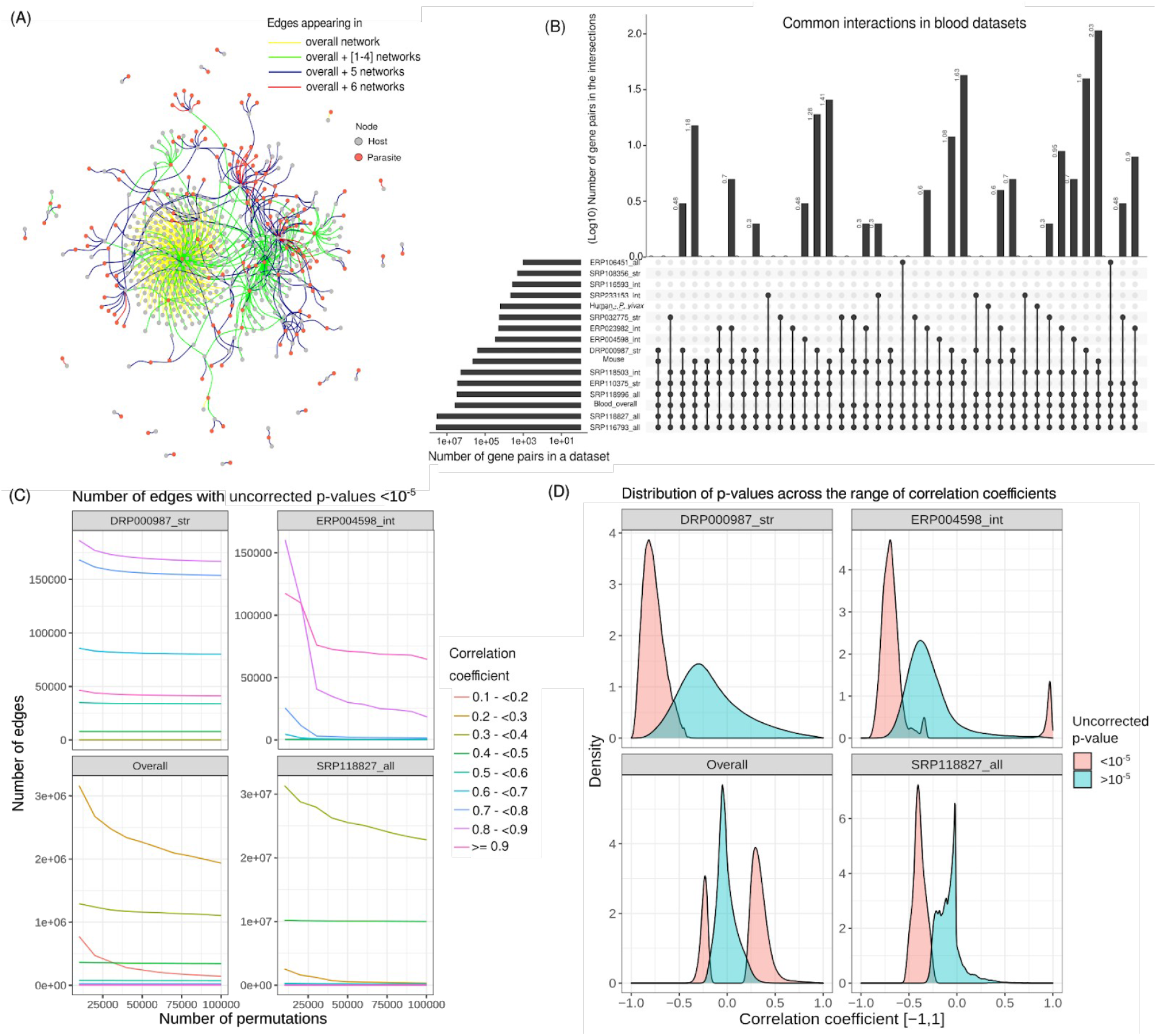
Multi-layer networks of host-parasite interactions across host-parasite systems increase resolution over neworks from single studies. We performed pairwise gene correlation tests on individual studies for gene expression in malaria and on an “overall” dataset constructed by concatenating data from 15 datasets. (A) A small part of the “overall” network shows those overlaps. We selected this part of the network starting from a set of “core edges” found in most networks (the “overall” network and networks from six individual studies). Edges in the neighbourhood to this were randomly selected in equal numbers, where available, based on how many networks, out of 15, they were found in. The number of networks an edge is found in is represented by the edge colour. Several edges were found common to multiple datasets across host-parasite systems, a result later used to derive a “core” network. (B) The (log 10 transformed) number of edges in networks from individual studies (horizontal bars) and shared (vertical bars) among networks shows the large overlaps between individual studies but also large differences in the sizes for those overlaps. (C) shows the effect of performing an increasing number of permutations on the p-value of an edge within a single network (datasets ERP004598_all, SRP118827_int, DRP000987_str and “overall” as examples). As the number of permutations increases, the increase in resolution (decrease in the number of significantly correlated edges recovered) slows, warranting a different approach to increase resolution, such as the multi-layer analysis depicted in previous panels. (D) shows the distribution of p-values across the range of correlation coefficients. The same studies as in fig (C) are used. While individual datasets might represent different aspects of malaria infection and have a reduced number of interactions in common, we here show the benefit of analysing the “overall” dataset: the “overall” analysis identifies both positive and negative correlations. In the results of the “overall” dataset, we reach a consensus of all the datasets included.

#### A “core” network of evolutionarily conserved interactions

The “overall” network is a highly connected graph with a total of ∼3.64 million edges (Supplementary Table S2A). Even though this network provides some resolution relative to the ∼56 million edges possible in a network of 13986 host and 4005 parasite genes, the resolution of this network might still be improved. We therefore defined a “core” network: using the “overall” network as a scaffold, we extracted edges that were recovered in at least one human study and at least one model organism. Using this definition, the resultant “core” network has 1876 host genes and 2050 parasite genes connected by 15324 edges (Supplementary Table S2B). A list of GO terms enriched or depleted in the genes of the “core” network are provided in Supplementary Table S3. As expected, many GO terms were the same as in the “overall” network. Most GO terms in the “overall” network were enriched more strongly because of the higher number of genes in that network. However, many GO terms in the “overall” network were broader than in the “core” network. The recovery of more specific functions in the “core” network indicates higher resolution in this network (Table S3).

#### Gene co-expression explains gene essentiality

A node is important in a network if it is central, i.e., highly connected to other nodes in the network. The metrics node degree (DG), betweenness (BW) and eigenvector centrality (EC) are network centrality measures that can quantify this. Similarly, an essential gene is classically described as a gene performing a function necessary for the viability of a cell or organism. We thus hypothesized central nodes in our parasite networks would tend to be more essential genes of *Plasmodium*.

In quantitative terms, more essential genes are more important for cell growth and thus, the disruption of those genes leads to larger growth defects. Such data is available for *P. berghei* in mice, for which [35] reported “relative growth rates” (RGR) of mutant parasites reflecting the essentiality of a gene. A second study [36] reported *P. falciparum* growth rates impacted by genome wide mutagenesis covering 5,399 protein-coding genes. Here, the Mutagenesis Index Score (MIS) quantifies essentiality of a gene.

First, we tested the different network metrics for predicting gene essentiality (RGR and MIS) as response variables in beta-regression models. EC generally resulted in the best models and combination of the centrality measures did not improve the models in most cases (Supplementary Table S4). This might be explained by DG and EC being tightly correlated. We therefore report models with EC as a single predictor.

We then compared how well EC from different networks explained gene essentiality (Table 3, Fig 5): EC in the network derived from human - *P. falciparum* studies did significantly predict *P. falciparum* MIS and *P. berghei* RGR. EC from *P. berghei* networks, was also a significant predictor for both *P. falciparum* MIS and *P. berghei* RGR. Surprisingly, *P. berghei* network centrality was a better predictor of MIS in *P. falciparum* than the centrality in *P. falciparum* expression networks. EC from both the “overall” network and the “core” network was a significant predictor for gene essentiality. Considering effect sizes, p-values and AIC, we conclude that centrality in the *P. berghei* network best explains both RGR and MIS, followed by the metric from the “core” network. Given the much smaller size of the “core” network, the predictive power of centrality within this network is striking.

**Table 3.**
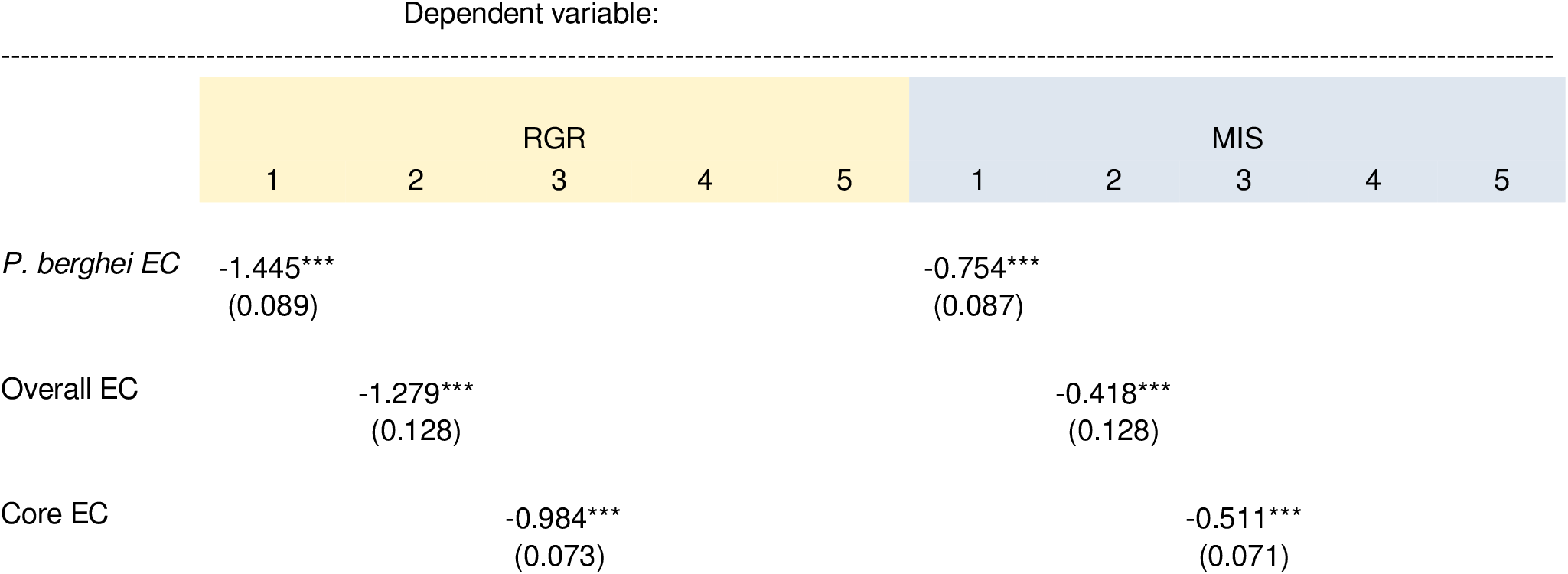
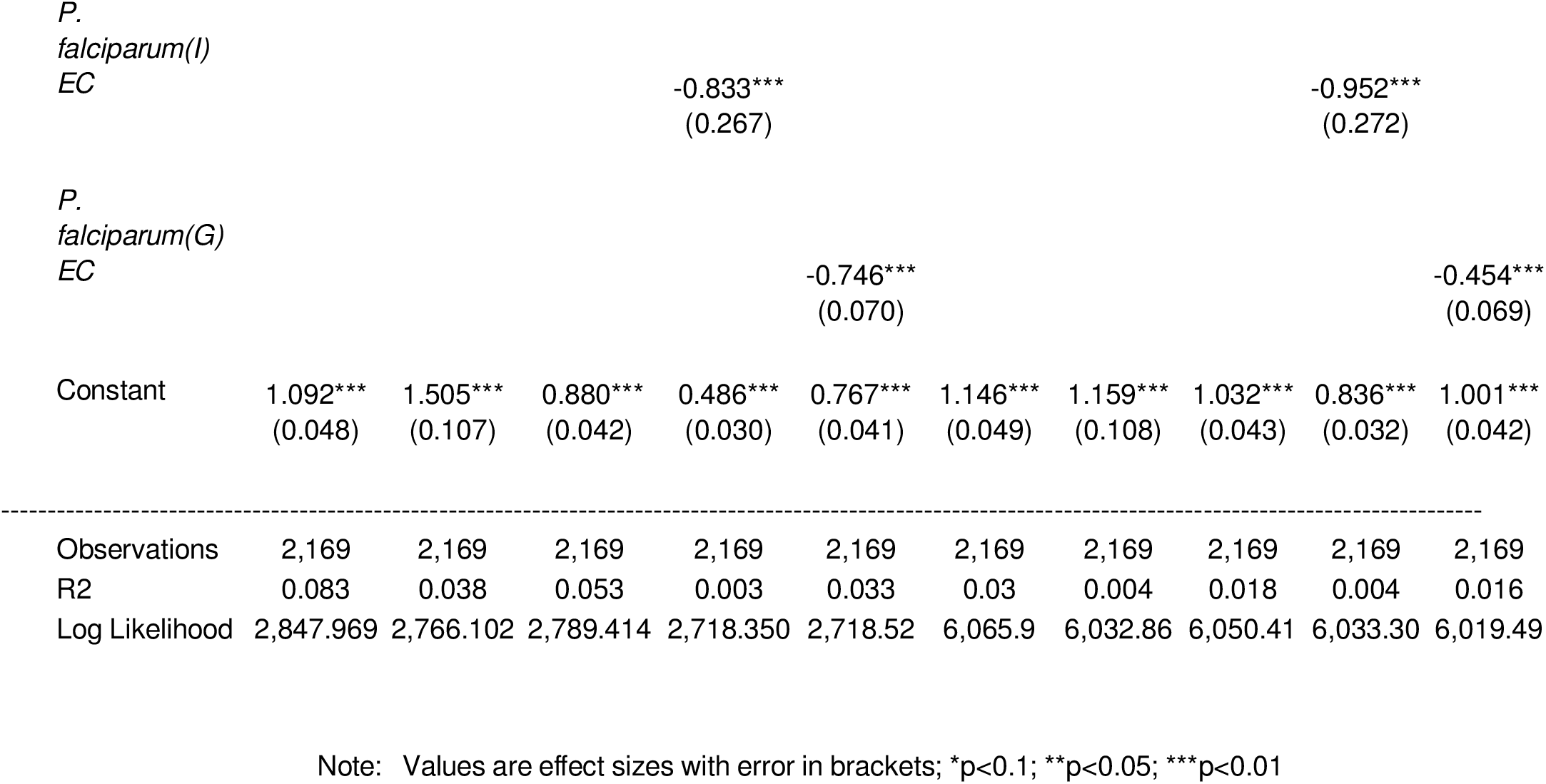
Overview of beta regression models to explain Relative Growth rate (RGR) and Mutagenesis Index Score (MIS) using eigenvector centrality measures of 5 networks: from 1) a single study on *P. berghei* infection in mice, from 2) overall, 3) core, 4) a single study on human *P*.*falciparum* infection in Indonesia (I) and 5) another singly study on human *P. falciparum* infection in Gambia (G).

**Figure 5.**
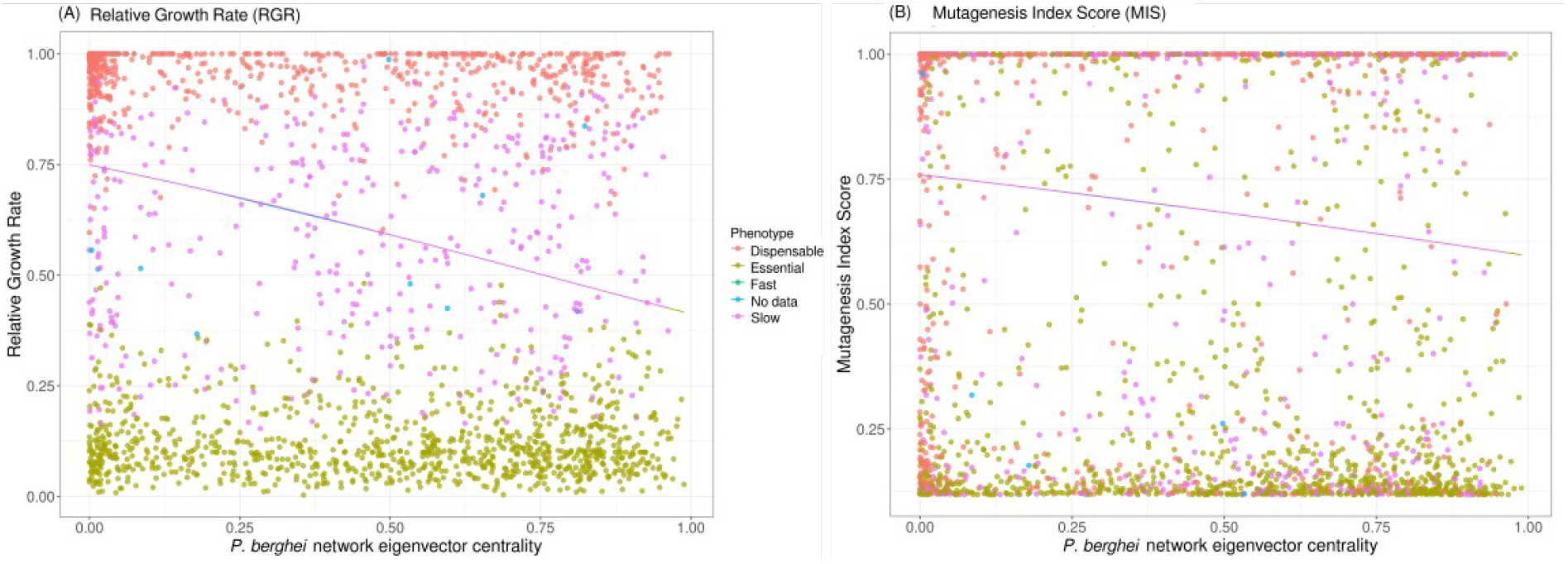
Explanation of gene essentiality with gene co-expression networks. Eigenvector centrality in a “core” network of correlated parasite-parasite gene expression is plotted against Relative Growth Rate (RGR, in A) and Mutagenesis Index Score (MIS, in B), respectively. Lines depict predictions of the essentiality scores from the centrality measure in a beta-regression: the higher the centrality, the lower is the essentiality score (RGR and MIS), meaning that genes central in the network are more essential to the parasites growth and survival. Color for both figures indicates phenotypes as categorised in [35].

We can conclude from this analysis, that a) our co-expression networks capture biological characteristics independently measured for the parasite and b) that parasite genes with a central position in the “core” network are important for the parasite’s growth and survival. Highly connected genes in the intra-parasitic “core” network, however, might control the reaction to host signals without necessarily directly interacting with host genes.

#### Interacting parasite processes and host immune response

Based on functional annotation of our networks, we show that the correlation between host and parasite transcript expression can highlight known processes important in host-parasite interactions (Fig 6, Table S3). Biological processes for hosts consistent among almost all networks include “Cell adhesion by cadherin” and “Calcium-dependent cell adhesion”. Cadherins are cell-adhesion proteins that depend on calcium. *Plasmodium* infection causes systemic endothelial activation of host blood vessels when the infected RBCs (iRBCs) sequester on the lining of blood vessels. This is accompanied with an increased interaction of endothelial cells with WBCs, as cell adhesion molecules on the vessel lining direct WBC trafficking to infected RBC sequestration sites [104]. In addition, immune cells like B cells and monocytes express adhesion-related genes found in this GO term, like Fer (Tyrosine-protein kinase) and ICAM-1 (Intercellular adhesion molecule 1), respectively [105]-[107]. To connect co-occurring host and parasite biological processes, we found associations between enriched (p-value <= 0.05) host and parasite GO terms based on the interactors of the genes in these enriched GO terms. From the “core” network, we identified a GO network of 617 host GO terms and 464 parasite GO terms. This analysis suggests that besides the detailed co-expression of gene products, broader enriched host and parasite processes or pathways are likely interacting in malaria infection (Fig 7; Supplementary Table 5).

**Figure 6.**
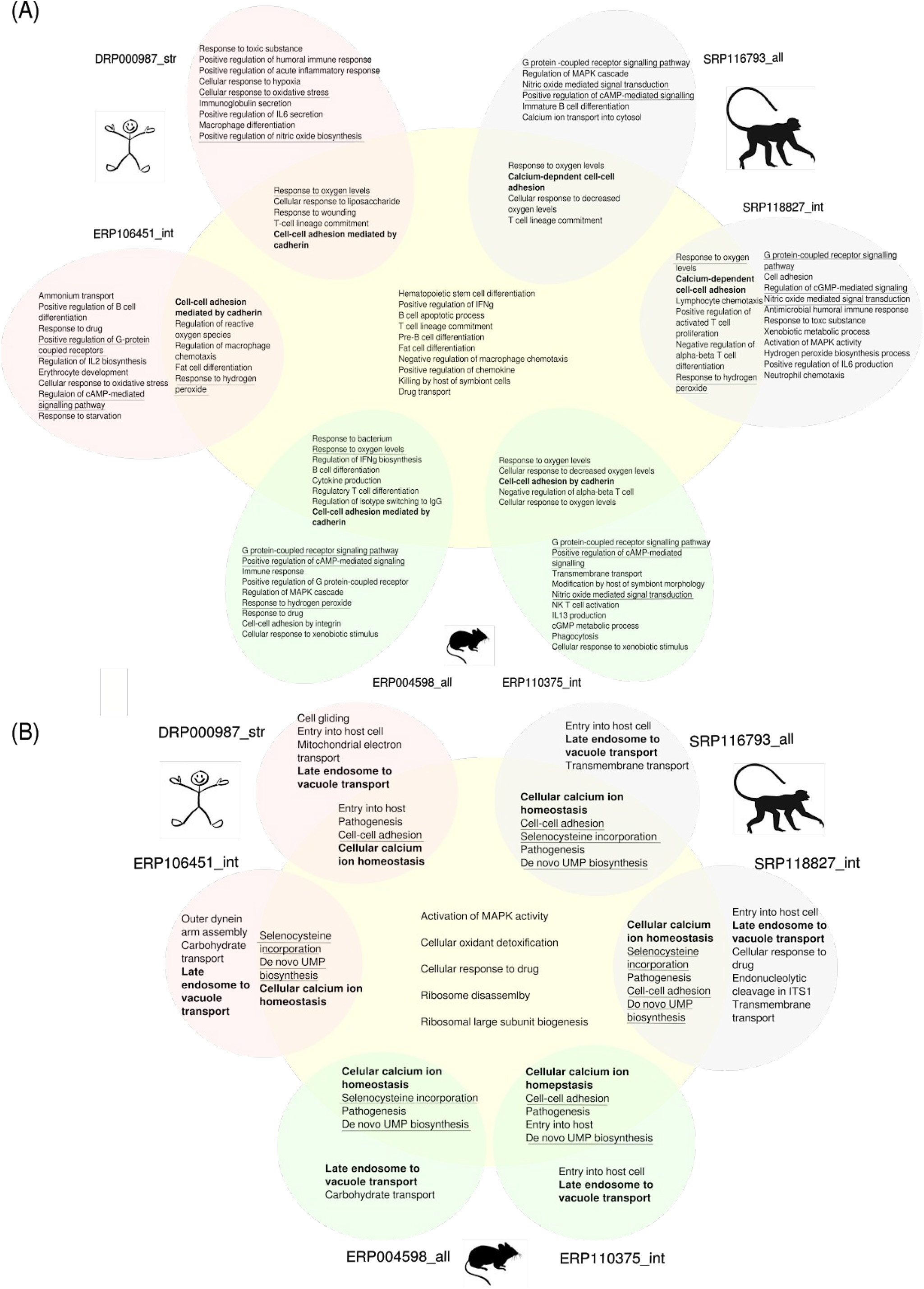
Biological Processes (BP) Gene Ontology (GO) terms shared between datasets across host-parasite systems. We performed GO term enrichment analysis for the “overall” dataset and its 15 constituent datasets for (A) host and (B) parasite genes. Here we show GO terms for the 6 largest studies for each host species. Enriched GO terms (p-value >= 0.05) are presented in sets coloured in red, green and blue for human, mouse and macaque studies. The GO terms of the “overall” dataset are displayed in the central yellow set. GO terms for the six individual datasets with the highest contribution to the “overall” dataset are shown in the perimeter of this. Overlapping between the respective areas shows shared GO terms found in either dataset. Set overlaps are not illustrated between the six individual datasets but respective terms are underlined (shared between 4-5 studies) or bold (shared between all 6 studies).

**Figure 7.**
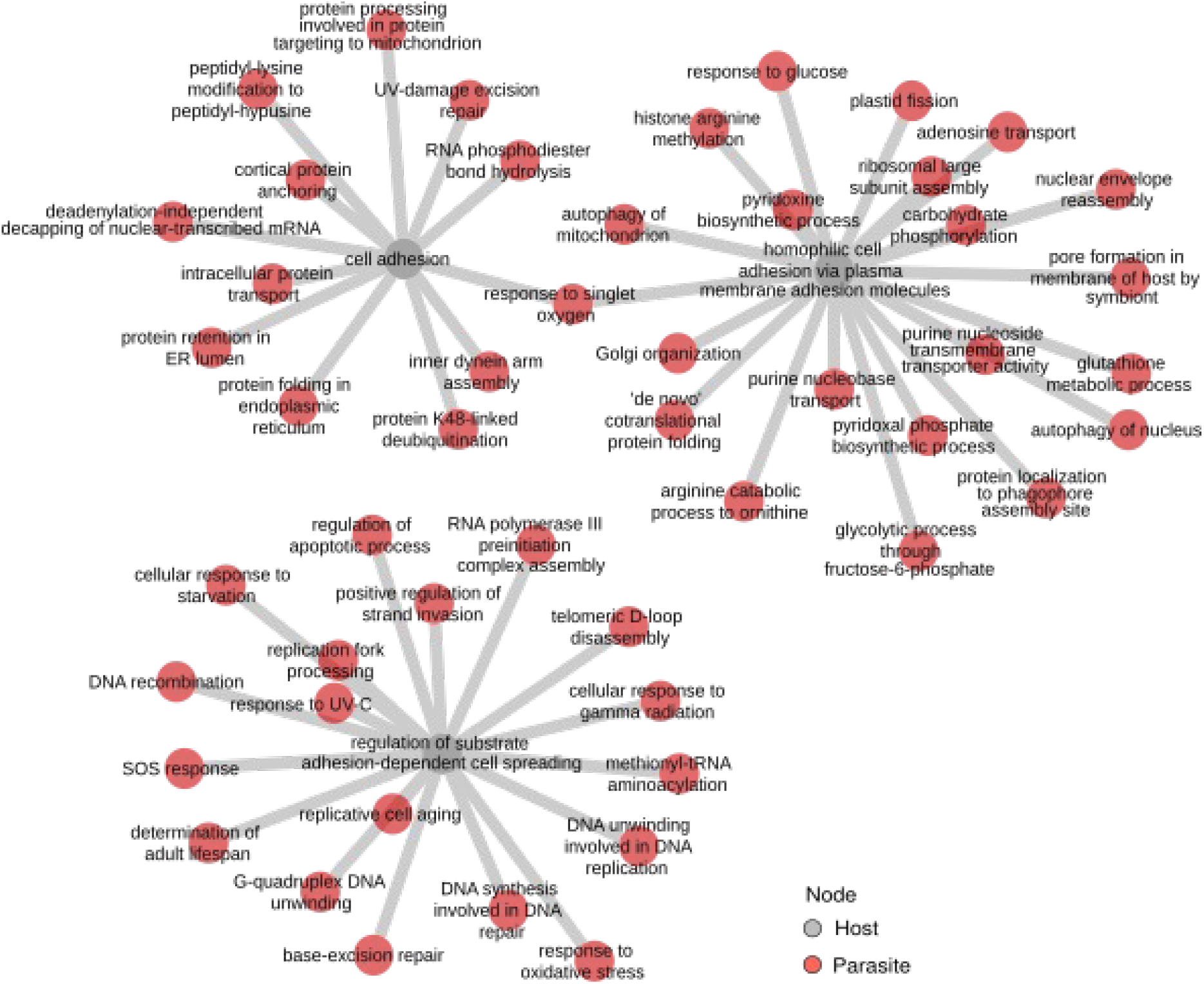
An interaction network of host-parasite GO terms. Host and parasite genes in the “overall” and “core” gene expression correlation networks were analysed for GO term enrichment. For each individual enriched host GO term interacting genes from the other species were then analysed for enrichment more specifically. This gives and understanding of what GO biological processes in one organism, say, the host, are associated with specific processes in the parasite. In this figure, we show all parasite GO terms associated with host GO terms related to “adhesion”, in the “core” network.

Another host biological process commonly found in our interaction networks is the response to oxidative stress combined with response to hydrogen peroxide and nitric oxide mediated signalling.

It has been found that iRBCs produce twice as many free OH^-^ radicals and H2O2 than uninfected RBCs [91,108]. This mechanism is believed to be a defence mechanism to abate the infection, although the exact mechanism of parasite killing is still unclear. Nitric oxide (NO) production by the host is also believed to promote parasite death in malaria [109]. An interesting GO term observed in the results was “Killing by host of symbiont cells”. This term included genes that were all involved in defence response towards microbes by invoking immune cells (eg, Neutrophil cytosol factor 1 and Cathepsin G), respiratory burst (Ncf1) and blood coagulation (eg, prothrombin).

Host-parasite interaction networks from all infection systems were consistently enriched for the parasite processes, “Calcium ion homeostasis” and “Late endosome to vacuole transport”. Calcium ion homeostasis regulated by calmodulins and Ca^2+^-dependent kinases (CDPKs) play roles in complex signalling pathways and are important for apicomplexan parasite virulence [110,111]. Vacuolar protein sorting-associated proteins-1, -2 and -46 (VPS1, 2 and 46) along with serine/threonine protein kinase (VPS15) are the underlying signal for the enrichment of “Late endosome to vacuole transport” process and are interlinked with host immune reaction genes in our interaction networks. The endomembrane system is important for the parasite in order to invade the host cell, to establish the parasitophorous vacuole and to obtain nutrients. The Endosomal Sorting Complex Required for Transport (ESCRT) is involved in late endosome formation as endosomes exists at different stages of formation - from early to late endosome, before finally fusing with the lysosome or vacuole (reviewed in [112]). The interrelation of the immune system and endosomal formation and sorting events should be a future focus of research including our deeper studies of correlated gene expression networks.

To test whether our networks (gene clusters) are indicative of correlated gene expression originating from specific immune cells, we used immune cell marker genes established in [113]. These marker genes are specific for 6 types of immune cells in the event of a *Plasmodium* infection in the blood: neutrophils and monocytes (innate immunity), T cells and B cells (adaptive immunity), and myeloid dendritic cells and NK cells (innate and adaptive). We found (Supplementary Table S6) that specific markers from neutrophils were significantly overrepresented (Fisher’s exact test, p-value = 0.0089) in our “overall” network. Specific markers from monocytes were not significant but overrepresented (Fisher’s exact test, p-value = 0.19). This likely indicates phagocytosis and killing of *Plasmodium* via respiratory burst in which neutrophils and monocytes have a central role [114,115]. Specific markers for myeloid dendritic cells had significant underrepresentation (Fisher’s exact test, p-value = 0.0027). Marker genes for all other cell types were underrepresented (Fisher’s exact test, p-value > 0.05). In the “core” network, marker genes were not significantly enriched, but those specific for monocytes, neutrophils and NK cells were tentatively overrepresented. We then used the presence of these specific cell type markers to annotate network clusters with the cell-type indicated (Fig 8). This resulted in 68 clusters in the “overall” network and 14 clusters in the “core” network annotated with a single cell type and additional 49 and 5 clusters annotated with multiple cell types, respectively. Among the most densely interconnected cluster in the “core” network with strong evidence of specific cell types was a set of 203 host genes (three are specific for T cells, two for neutrophils) correlating in their expression with only two parasite genes. One of them, a ubiquitin regulatory protein (PBANKA_1222400), is connected to 269 host genes in total in the “core” network. It appears to be uniformly expressed in all stages of the *Plasmodium* life cycle [34]. WLL-vs (Leu-Leu-Trp vinyl sulfone) and WLW-vs (Trp-Leu-Trp vinyl sulfone) are proteasome inhibitors and likely drug candidates [116]. In an experiment to study genetic changes mediating parasite recrudescence in WLL-vs and WLW-vs resistant mutants, this protein’s level was amplified in recrudescent lines [117]. Thus it was suggested to confer low-grade resistance to such proteasome inhibitors. The second *Plasmodium* gene is a putative dynamin-like protein (DrpC, PBANKA_1434100), connected to 11 host genes. Dynamins are mechanochemical enzymes with a GTPase domain and one of the functions of dynamin-related proteins (Drp) is to traffic vesicles [118]. DrpC is conserved in apicomplexans and was shown to be localized at the base and periphery of the mitochondrion in *Toxoplasma gondii*, implying a probable role in mitochondrial fission [119]. Another module of three host genes included a NK cell marker correlated with two parasite genes - a 26S proteasome subunit (PBANKA_1206600) and ubiquitin fusion degradation protein UFD1 (PBANKA_1024700). The parasite 26S protease subunit [120]-[122] was reported as an essential protein in *P. berghei* but not in *P. falciparum* [35,36]. In general, the ubiquitin-proteasome system (UPS) is essential for quality control of proteins in eukaryotes. In *Plasmodium*, the UPS is expressed across all life cycle stages and is speculated to be a promising drug target. It helps adapt the parasite to stress, such as changes in oxidative environment and temperature differences, ensuring the survival and virulence of the parasite. The *P. falciparum* ortholog of UFD1 (PF3D7_1418000) was reported to be a pathogenesis-related protein in an *in silico* module-based subnetwork alignment approach using protein-protein interactions of *E. coli* as references [123].

**Figure 8.**
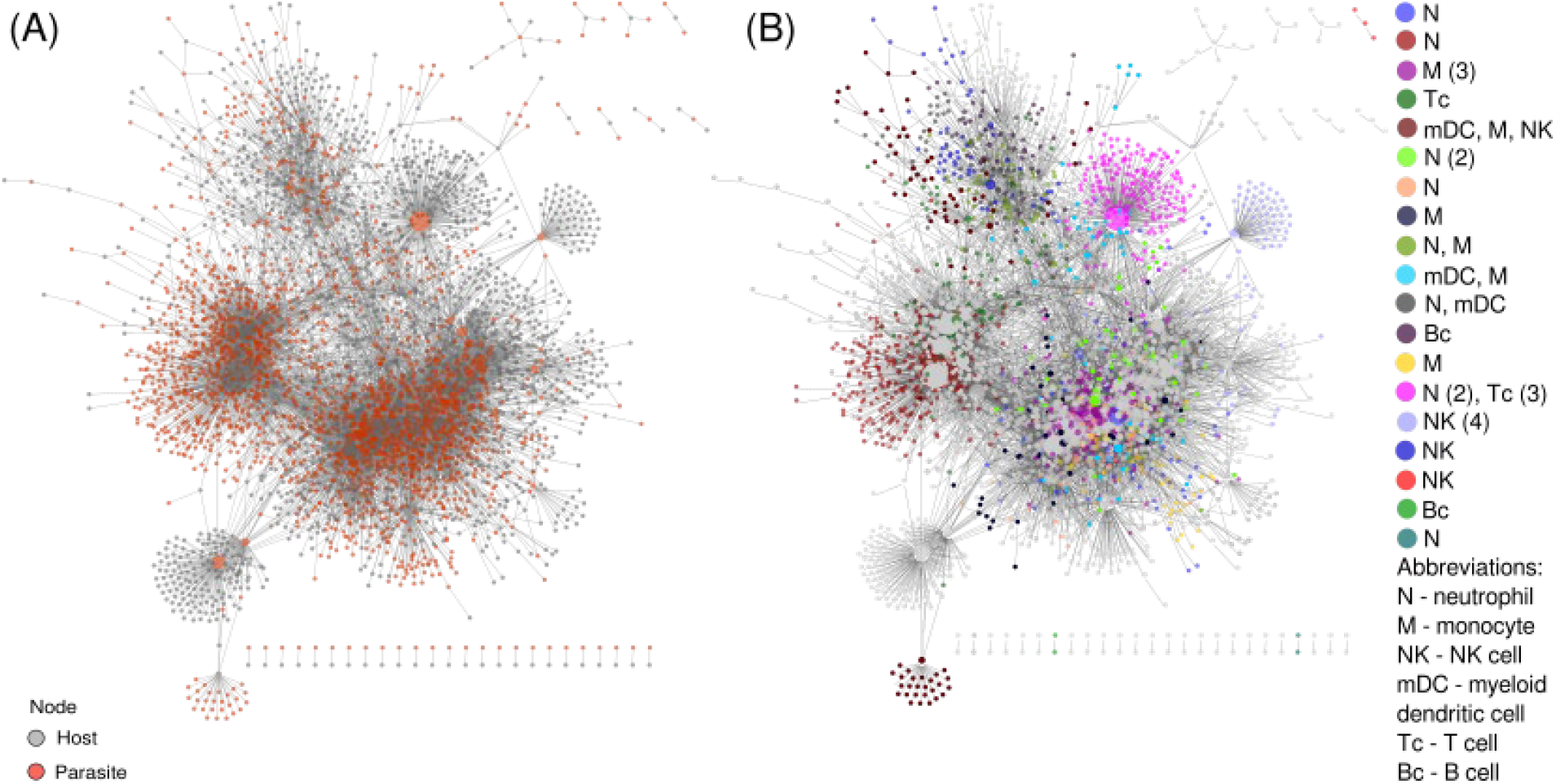
A “core” network highlights parasite interaction with specific immune cells. We derived a “core” network containing interacting genes supported by correlated gene expression in at least one human and one model organism study, A) this complete “core” network is displayed with grey nodes for host and red nodes for parasite genes. The size of a node is based on its connectivity - the higher its degree, the larger is the node. (B) illustrates the identification of node clusters in the “core” network using an edge betweenness algorithm from igraph. Clusters with immune cells specific marker genes are coloured (see legend, including a count for the number of marker genes in brackets). The remaining clusters are left grey.

One module had genes specific to three different immune cells - monocytes, myeloid DC and NK cells - the highest number of distinct immune cell types represented in a “core” network cluster. Among the 46 co-expressed parasite genes in this module were genes related to apicoplast biogenesis, mitochondrial fission, transcription, vacuolar transport from Golgi apparatus to the endoplasmic reticulum and Fe-S cluster assembly proteins.

Overall, these results (Supplementary Table S6) give an indication that certain gene expression clusters in our networks might be associated with specific cells of the innate immune response (neutrophils and monocytes). In addition, we show that parasite processes likely invoking this immune response include the expression of genes involved in drug resistance and in vesicular transport.

We next looked at overlaps between networks in our multilayer network analysis. Twenty interactions of specific gene pairs were conserved across 6 individual networks and the “overall” network (Fig 4A): this includes i) negatively correlated (Pearson’s rho = -0.26) Kelch13 in *Plasmodium* with Laminin subunit beta-2 (LAMB2) in the host and ii) negatively correlated (Pearson’s rho = -0.33) parasite 26S protease subunit and host LAMB2, both recovered only in mouse and monkey studies. LAMB2 is an extracellular high-affinity receptor and is associated with GO term “substrate adhesion-dependent cell spreading” to which infected erythrocytes (IE) have been reported to bind, along with to other adhesion molecules such as ICAM, VECAM, etc. [124, 125] found that laminin levels in serum were increased in severe malaria. [126] later contradicted this, finding that the binding of infected erythrocytes to endothelial receptors including laminin was the same in severe and non-severe malaria. Kelch13 is a well-studied protein in which a C580Y mutation in *P. falciparum* confers resistance to artemisinin, a frontline anti-malarial drug. It was found to be essential in both *P. falciparum* and *P. berghei* intraerythrocytic stages. Associated proteins and Kelch13 form an endocytic compartment and are essential for feeding on host hemoglobin. Kelch13 is also suggested to be a ubiquitin ligase with a role in the ubiquitin-proteasome system by labeling proteins for degradation [127]. Artemisinin and its derivatives (ART) are known to be activated by the products of hemoglobin degradation and promote cell death by increasing ER stress facilitated by the accumulation of polyubiquitinated proteins. Resistance-conferring mutations on Kelch13 reduce host cell and hemoglobin endocytosis and along with 26S proteasome system (which includes the 26S protease subunit, our other parasite gene in discussion here), maintain the normal proteasomal degradation pathway preventing cell death, and thus, resulting in resistance [128-132]. Our work here is, to our knowledge, the first suggestion that Kelch13 and LAMB2 might be involved in interacting host-parasite processes.

The gene pairs discussed above were recovered in 7 datasets altogether (6 individual + “overall”), but were not recovered by any of the human - *Plasmodium* studies and are therefore not present in the “core” network. Of the 20 interactions that were found in 7 datasets, 5 were also found in the “core” network and are indicated as such in the Supplementary Table 2C. Of these 5, one was an interaction between host protein Odorant-receptor ODR4 homolog (ENSG00000157181) and parasite Thioredoxin-like associated protein 2 (TLAP2; PBANKA_0518100). TLAP2 is a microtubule-associated complex. It was found to be dispensable in both *P. berghei* and *P. falciparum* [35,36]. TLAP2 is conserved in *Toxoplasma gondii* and *Plasmodium* spp. [133]. Along with other TLAP proteins, it is associated with protein TrxL1 (Thioredoxin-Like protein 1) and as a complex, coats cortical microtubules [133]. These coating proteins, as an ensemble, stabilize the cortical microtubules [134]. Host ODR4 is involved in trafficking of GPCR proteins [135] and in protein localisation [136].

A second interaction was between host biotinidase (ENSG00000169814) and parasite with protein kinase (PBANKA_1016200). In the host, biotinidase makes biotin available from dietary sources. Host biotin was found to be essential for *Plasmodium* survival in the liver stages but not in the blood stages [137]. In general, several mutations in biotinidase reduce biotin availability without causing severe diseases. There have been suggestions that biotinidase mutations could be used as antimalarial therapies, following evidence that there are biotinidase mutations in blood spots of Somali populations [138]. Putative parasite protein kinase PBANKA_1016200 is a membrane protein involved in protein phosphorylation and ATP binding. Even though it is dispensable in both *P. berghei* and *P. falciparum*, in general, protein kinases have been an attractive subset of proteins to be used as antimalarial targets (reviewed in [138]).

We propose that interactions between these proteins (physical protein-protein interaction) or interlinkage of associated pathways may be worth further scrutiny in mechanistic investigations.

## Conclusion

Our analysis recovers both very broad and well-known processes involved in host-parasite interactions as well as strong interaction signals for narrower processes, including specific interacting gene pairs. This indicates that our analysis of interaction networks might uncover novel links between host and parasite processes helping in focus further research into detailed mechanisms of host-parasite interactions. Further analyses building on the methodology and concepts conceived here will involve networks derived from liver studies curated here, in which host and parasite cells interact more directly than in the blood. Methodologically, we show that analyses of expression from different host-parasite systems as multilayer networks can link host and parasite network modules. Meta-analysis of co-expression networks can thus assess whether host-parasite interactions are shared among different studies and treatments, different tissues and different host-parasite systems. This allows the identification of processes at different scales of molecular pathway organisation as depicted above.

Moreover, the sharing of pathways among co-expression networks from different host-parasite systems might indicate how easily results on those pathways can be translated between different host-parasite systems - here presented in a “core” network. Our results can thus provide insights into how easily and for which pathways observations made in malaria models can be translated to human malaria.

## Supporting information

Malaria_DualRNAseq_SupplementaryInfo

## Conflict of Interest

The authors declare that there is no conflict of interest.

## Author Contribution

EH and PM designed the study. PM obtained and analysed the data. EH, GB and PM wrote the manuscript. All authors contributed to the final version of the manuscript.

## Acknowledgement

This research project was undertaken with the assistance of resources and services from the National Computational Infrastructure (NCI), which is supported by the Australian Government. We thank the statistics user group of the Leibniz-IZW for very helpful comments on our analysis.

## Funding

This work was supported by the Alliance Berlin Canberra “Crossing Boundaries: Molecular Interactions in Malaria”, which is co-funded by a grant from the Deutsche Forschungsgemeinschaft (DFG) for the International Research Training Group (IRTG) 2290 and the Australian National University. G.B.’s work is supported by the National Health and Medical Research Council of Australia (Grant APP1143008), the Australian Research Council (Grant DP180101494) and the National Natural Science Foundation of China (Grant 81772214)

## Notes

### Competing Interest Statement

The authors have declared no competing interest.

### Summary of Updates

Major revisions: A core network is defined; Explanation of intra-parasitic gene co-expression network using gene essentiality (external data) added; Identification of immune cell marker genes (external data) in network clusters added; Figures updates, more supplementary files added

